# SARNAclust: Semi-automatic detection of RNA protein binding motifs from immunoprecipitation data

**DOI:** 10.1101/127878

**Authors:** Ivan Dotu, Scott Adamson, Benjamin Coleman, Cyril Fournier, Emma Ricart-Altimiras, Eduardo Eyras, Jeffrey H. Chuang

## Abstract

RNA-protein binding is critical to gene regulation, controlling fundamental processes including splicing, translation, localization and stability, and aberrant RNA-protein interactions are known to play a role in a wide variety of diseases. However, molecular understanding of RNA-protein interactions remains limited, and in particular identification of the RNA motifs that bind proteins has long been a difficult problem. To address this challenge, we have developed a novel semi-automatic algorithm, SARNAclust, to computationally identify combined structure/sequence motifs from immunoprecipitation data. SARNAclust is, to our knowledge, the first unsupervised method that can identify RNA motifs at full structural resolution while also being able to simultaneously deconvolve multiple motifs. SARNAclust makes use of a graph kernel to evaluate similarity between sequence/structure objects, and provides the ability to isolate the impact of specific features through the bulge graph formalism. SARNAclust includes a key method for predicting RNA secondary structure at CLIP peaks, RNApeakFold, which we have verified to be effective on synthetic motif data. We applied SARNAclust to 30 ENCODE eCLIP datasets, identifying known motifs and novel predictions. Notably, we predicted a new motif for the protein ILF3 similar to that for the splicing factor hnRNPC, providing evidence for interaction between these two proteins. To validate our predictions, we performed a directed RNA bind-n-seq assay for two proteins: ILF3 and SLBP, in each case revealing the effectiveness of SARNAclust in predicting RNA sequence and structure elements important to protein binding. Availability: https://github.com/idotu/SARNAclust

## Background

RNA-protein binding is a fundamental biological interaction vital to the diverse functions of RNA, including key roles in RNA splicing, translation, localization and stability (Hafner et al. 2010; Lukong et al. 2008; Sanford et al. 2009; Yeo et al. 2009). However, the sequence features that determine affinity to RNA-binding proteins (RBPs) are unknown for most RBPs, including the vast majority of the hundreds of RBPs in the human proteome. Moreover, even for RBPs with known binding motifs, existing sequence motifs are only weakly predictive of which RNA regions will be bound. Deciphering these RNA binding features is crucial for mechanistic understanding of RNA-protein binding. The development of quantitative models that predict RBP targets, the strength of binding, and the sequence regions that control such binding will be paramount for understanding how RNA regulation impacts human health. RNA-protein interactions are already known to play a role in a wide variety of diseases including muscular dystrophy, fragile X syndrome, mental retardation, Prader-Willi syndrome, retinitis pigmentosa, spinal muscular atrophy, and cancer (Hafner et al. 2010; Lukong et al. 2008; Sanford et al. 2009; Wurth 2012; Yeo et al. 2009).

To date, the RNA features that determine the binding of an individual protein have been studied primarily from the perspective of short individual motifs. For example, the 5 nt motif GGAGA is considered the canonical description for the RNA motif interacting with the human LIN28 protein (Wilbert et al. 2012, 28). In the RBPDB database (Cook et al. 2011) which compiles RNA-protein interaction motifs based on previous experimental findings, this type of short single motif is the standard descriptor for a binding element. Unfortunately, such short nucleotide motifs for RNA-protein interactions have often had poor predictive power. As an example, Hogan et al identified transcripts bound to 40 yeast RBPs, and then searched UTR regions of these transcripts for overrepresented motifs (Hogan et al. 2008). They were able to find statistically significant motifs for only 21 RBPs, and in many cases previously known motifs could not be found. This issue of poor predictive power for single motifs has continued even with finer resolution assays such as CLIP-seq, which can localize binding sites to within a few nucleotides.(Chi et al. 2009; Zhang and Darnell 2011). Wilbert et al. used CLIP-seq to find *LIN28*–RNA interaction sites in human somatic and embryonic stem cells (Wilbert et al. 2012, 28). Target transcripts were found to have 3.5 binding sites on average, with sites typically ∼35 nt or shorter and often >50 nt apart on transcripts. The GGAGA sequence motif was the most overrepresented motif in these sites, yet less than 13% of the sites contained it.

Computational approaches for RNA motif detection have had moderate success. RNA motif analysis has often been carried out by repurposing DNA motif finder tools such as MEME (Bailey et al. 2006), PhyloGibbs (Siddharthan et al. 2005) or cERMIT (Georgiev et al. 2010), but a fundamental limitation of these methods is that they cannot take into account RNA secondary structure. DNA-based tools have been partially successful because most known RBPs bind to single stranded RNA (ssRNA), but it remains unclear how much secondary structure impacts binding. Some motif identification methods have incorporated aspects of RNA structure, e.g. by biasing for single stranded regions (Hiller et al. 2006; Wang et al. 2011) or searching over a limited set of structural contexts (paired, loop, unstructured, miscellaneous) (Bahrami-Samani et al. 2015; Fukunaga et al. 2014; Kazan et al. 2010). However, the predictive power of these methods remains low, likely because of the limited number of considered contexts compared to the diversity of possible RNA structures. For example, Kazan et al. tested their algorithm on 9 RBP-interaction sets and found an average AUC value of only 0.64 (Kazan et al. 2010). More recently, new approaches that consider these structural contexts have arisen, using probabilistic machine learning algorithms such as Support Vector Machines (Livi and Blanzieri 2014), Hidden Markov Models (Weyn-Vanhentenryck and Zhang 2016; Zhang et al. 2013) or even Deep Learning (Alipanahi et al. 2015; Pan and Shen 2017; Zhang et al. 2016). Still, none of these latest methods are unsupervised (i.e. they rely on a training set of true binding sites), they do not consider multiple RNA motifs, and they abstract structural constraints rather than considering exact stable structures.

More relevantly, Maticzka and colleagues developed the graph kernel-based GraphProt and applied it to learn motifs from CLIP-seq data (Maticzka et al. 2014). They found motifs that were predictive of binding for the protein PTB, and the certainty of predicted motifs correlated with measured RBP affinities. However, the efficiency of this method is untested for RBPs that bind to double stranded RNAs, and it is unknown whether the effectiveness of the method would depend on the specific types of structures to which individual proteins bind. Moreover, GraphProt is not designed to deal with an RBP capable of binding several distinct RNA motifs. GraphProt reports at most one motif and classifies the remaining data as noise. A more general approach would be to use clustering to allow for multiple possible binding motifs. A related method is GraphClust (Heyne et al. 2012), which uses a sequence/structure graph kernel approach to cluster RNAs. However, it is tailored to cluster non-coding RNAs into families, and it is unknown if such an approach would be effective for the clustering of CLIP-seq sites.

Here, we propose a method, SARNAclust (Semi-Automatic RNA clustering), to cluster, as opposed to classify, RNA motifs that bind to a given RBP from CLIP-seq data. To the best of our knowledge, this is the first approach to attempt to cluster CLIP-seq peaks in order to discover potentially multiple distinct RNA motifs that bind to a given RBP. The most related approach we know of is AptaTrace (Dao et al. 2016), which uses clustering to identify multiple possible RNA motifs from HT-SELEX Experiments. However, AptaTrace cannot be applied to CLIP-seq sites since it relies on k-mer context information during evolution of a sequence pool over multiple SELEX rounds, while CLIP-seq provides a static snapshot. A key strength of SARNAclust is that it is fully unsupervised, an important feature since the noise levels in CLIP data are not well-characterized.

The paper is organized as follows: we first describe the algorithmic basis for the method and pipeline; we then perform a benchmarking on synthetic data and present results from analysis on 30 ENCODE datasets; next, we present experimental validations of the results using RNA bind-n-seq (Lambert et al. 2014) and gel shift assays; and, finally, we present a discussion and conclusions.

## Results

### Overview of the Algorithms

Two of the main results of this paper are: the development of a new RNA structure prediction method for CLIP peaks that we refer to as RNApeakFold; and the development of a new tool, SARNAclust, that includes RNApeakFold as part of a complete process to determine RNA motifs that bind to a given RBP. Our overall pipeline provides means to process data files coming from a CLIP experiment (see Methods) as well as source code for: (I) calculating potential secondary structure of the peaks using RNApeakFold, (II) clustering of peaks according to sequence only, and (III) clustering of peaks according to sequence/structure using SARNAclust. In addition, we provide a protocol for experimental validation of candidate motifs, including *in silico* design of instances of the motif using RNAiFold (Garcia-Martin et al. 2013, 2015). Figure 1 shows the flowchart of our pipeline.

**Figure 1.**
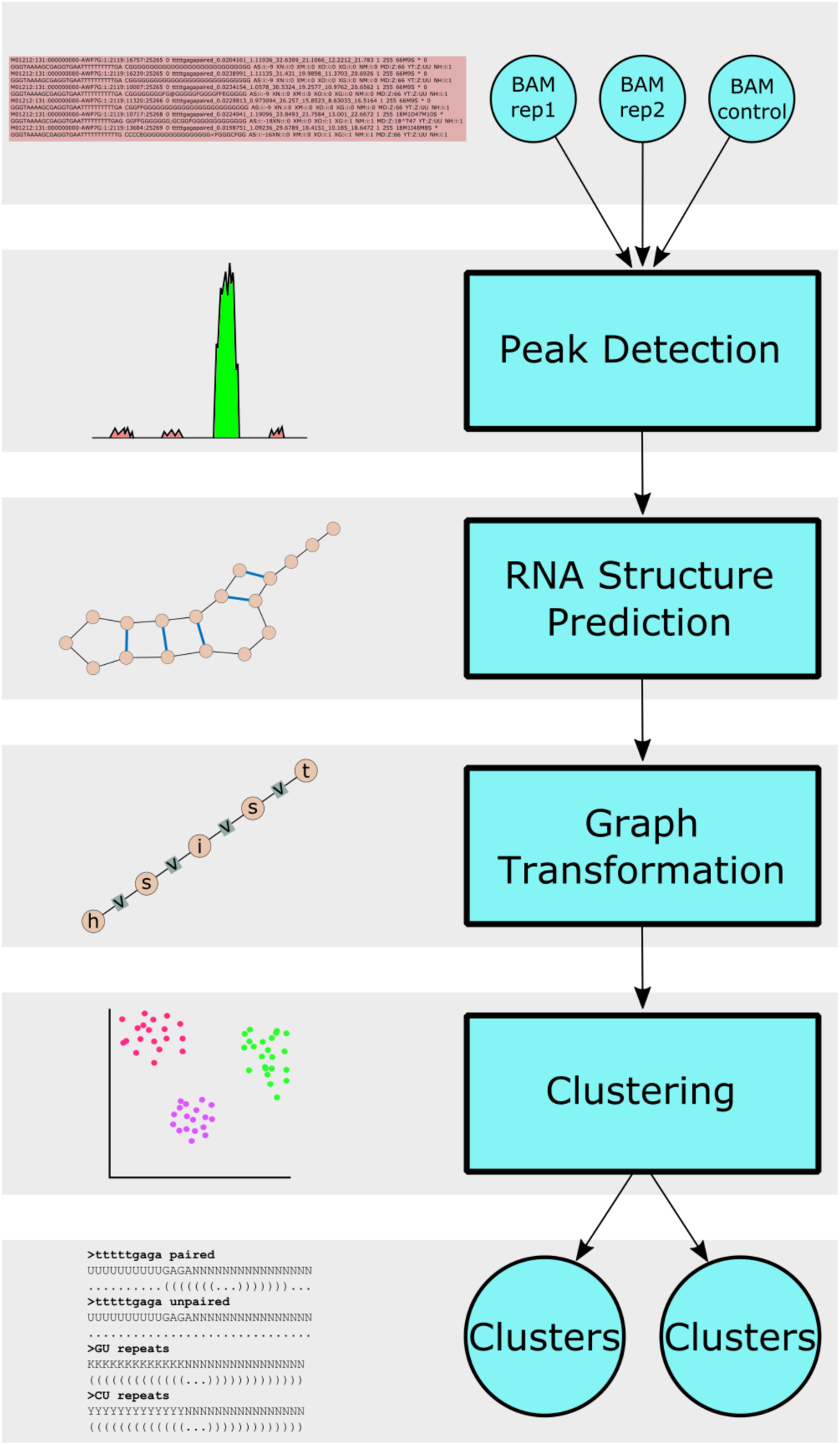
<u>Legend</u>: SARNAclust CLIPseq motif finding pipeline. Bam files for sample and control are processed through our peak detection module. The structure of each peak is calculated using RNApeakFold. The peaks are then transformed into sequence/structure objects and clustered˙.

Supplementary Figure 1 shows the flowchart of the peak analysis. Details of the experimental validation protocol can be found in the Methods section.

Broadly speaking, SARNAclust is a semi-automatic approach to find RNA motifs that bind to a given RBP. The key elements of SARNAclust are a) the structure calculation for CLIP peaks (RNApeakFold), and b) a graph transformation that allows for the calculation of a similarity value between pairs of sequence/structures. These similarity values provide the input for the clustering of CLIP peaks. Flexible parameters in SARNAclust allow it to be used as a guidance system to identify well-supported motifs and test their key features.

### RNApeakFold

To determine RNA secondary structure at each CLIP peak so that structure can be accounted for in motif detection, we have developed RNApeakFold, which calculates the probable secondary structure through computational folding. RNApeakFold first computes base pairing probabilities for the sequence including the peak and +/− 100 flanking nucleotides using RNAfold –p (Hofacker 2009). We then use those probabilities as energies in an implementation of Nussinov folding (Nussinov and Jacobson 1980) to determine the most probable structure of the peak region without the flanking nucleotides. As described below, this choice of folding approach yields superior motif detection. Algorithmic details are available in the Methods section.

### SARNAclust

Given the set of RNA sequence/structures calculated using RNApeakFold (or any other RNA structure prediction method), SARNAclust then clusters them. Similarities between pairs of sequence/structures are computed using the graph kernel in Eden (Costa and De Grave 2010), which is equivalent to that used in GraphClust (Heyne et al. 2012) and GraphProt (Maticzka et al. 2014). To use the graph kernel we first need to transform the sequence/structures into graphs. Our pipeline allows for several different transformations based on either the complete graph or the bulge graph (Kerpedjiev et al. 2015) (See Figure 2). The complete graph represents the secondary structure with all node connections between consecutive nucleotides and base pairs. The bulge graph is a condensed representation similar to the concept of abstract RNA shape (Giegerich et al. 2004). SARNAclust provides the following options for graph transformations (see Supplementary Figure 2):

- Option 1: GraphProt-like consists of the complete graph plus a hypergraph, which is a less condensed version of the bulge graph.
- Option 2: GraphProt-like where the hypergraph is substituted by the bulge graph.
- Option 3: Bulge graph.
- Option 4: Bulge graph plus corresponding sequence in hairpin loops.
- Option 5: Bulge graph plus corresponding sequence in internal loops and bulges.
- Option 6: Bulge graph plus corresponding sequence in external loops.
- Option 7: Bulge graph plus corresponding sequence in base paired regions.
- Option 8: Bulge graph plus corresponding sequence in hairpin loops, internal loops and bulges.
- Option 9: Bulge graph plus corresponding sequence in all unpaired regions.
- Option 10: Bulge graph plus sequence everywhere.
- Option 11: Bulge graph plus complete graph where sequence in base paired regions is not taken into account.

**Figure 2.**
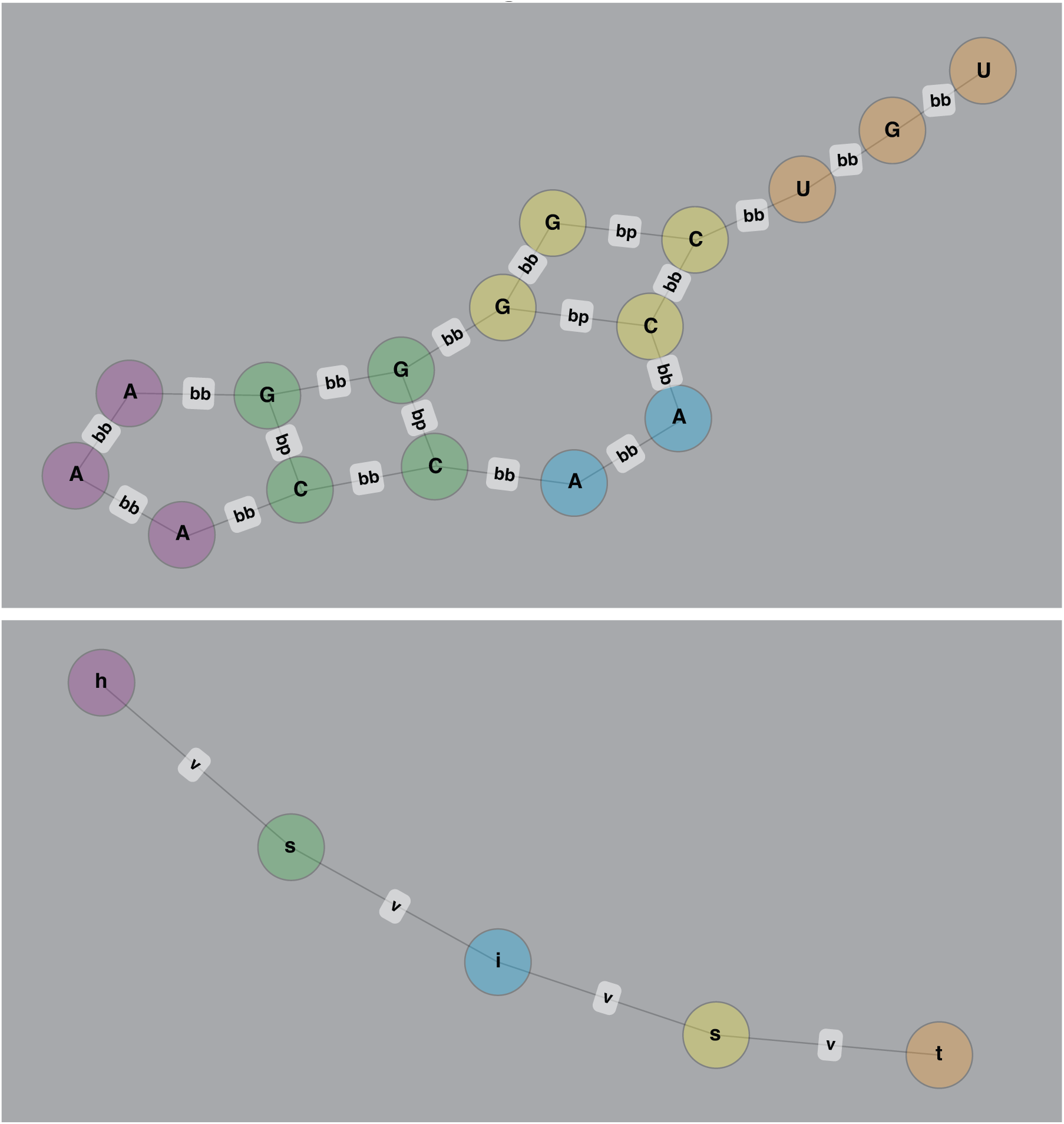
<u>Legend</u>: Complete graph and bulge graph sequence/structure representations used in SARNAclust. (top) complete graph and (bottom) bulge graph for example sequence/structure GGGGAAACCAACCUGU ((((…))˙.))… In the complete graph (top) nodes are nucleotides and edges between nodes correspond to either base pairing (bp) or backbone links (bb). In the bulge graph (bottom) the structure is collapsed into structural elements, where “h” is hairpin loop, “i” is internal loop or bulge and “s” is stem (double stranded). “t” stands for the three prime single stranded region

We have provided a range of options because different RNA-binding proteins will vary in their dependence on different structural features, and in many cases such features are known based on the domains in the protein. These include options that exhaustively consider structure but may be more sensitive to noise (e.g. option 1) and those that reduce the set of considered structural contexts based on prior expectations (e.g. option 11). For example, for options 9 or 11 to be suitable, one would hypothesize that the key binding element in the RNA tends to occur in unpaired regions, but within a precise structural context. In the following section we show the effect of these transformations when applying our methods to synthetic data.

Once the graph transformation has been applied, SARNAclust allows the user to apply one of several clustering algorithms and returns the clusters along with a consensus sequence/structure for each. The inputs needed to run the clustering module are: 1) the file with the sequence/structures, 2) dimension and 3) radius from Eden (see (Heyne et al. 2012), 4) the clustering algorithm and its parameters, and 5) the graph transformation option. Detailed description of the pipeline, manuals and source code are available at https://github.com/idotu/SARNAclust.

### Benchmarking on Synthetic Motif Data

In order to test the effectiveness of RNApeakFold and SARNAclust, we generated 100 sequences for each of the 6 synthetic motifs in table 1 plus 1000 random sequences to act as noise. Each synthetic motif describes the RNA motif that would bind a potential protein binding domain. The 6 motifs chosen correspond to: a special structure with no sequence conservation (*special_structure*) or a conserved sequence within a certain structural context in a hairpin loop (*GAGA_in_Hairpin*), in a bulge (*AUG_in_Bulge*), in an external loop (*pyrimidine_tract*) or in a double stranded region (*GGUCG_in_left_stem* and *GGUCG_in_right_stem*). Sequences for each motif were generated using RNAdualPF (Garcia-Martin et al. 2016) by sampling from the low energy ensemble of sequences compatible with the given structure and with the corresponding sequence constraints (see table 1). The 1000 random sequences were generated uniformly randomly (i.e, sampling each nucleotide with 0.25 probability) with a constant length comparable to the range of lengths of the synthetic motifs. All motif and random sequences can be found in Supplementary Data 1.

**Table 1:**
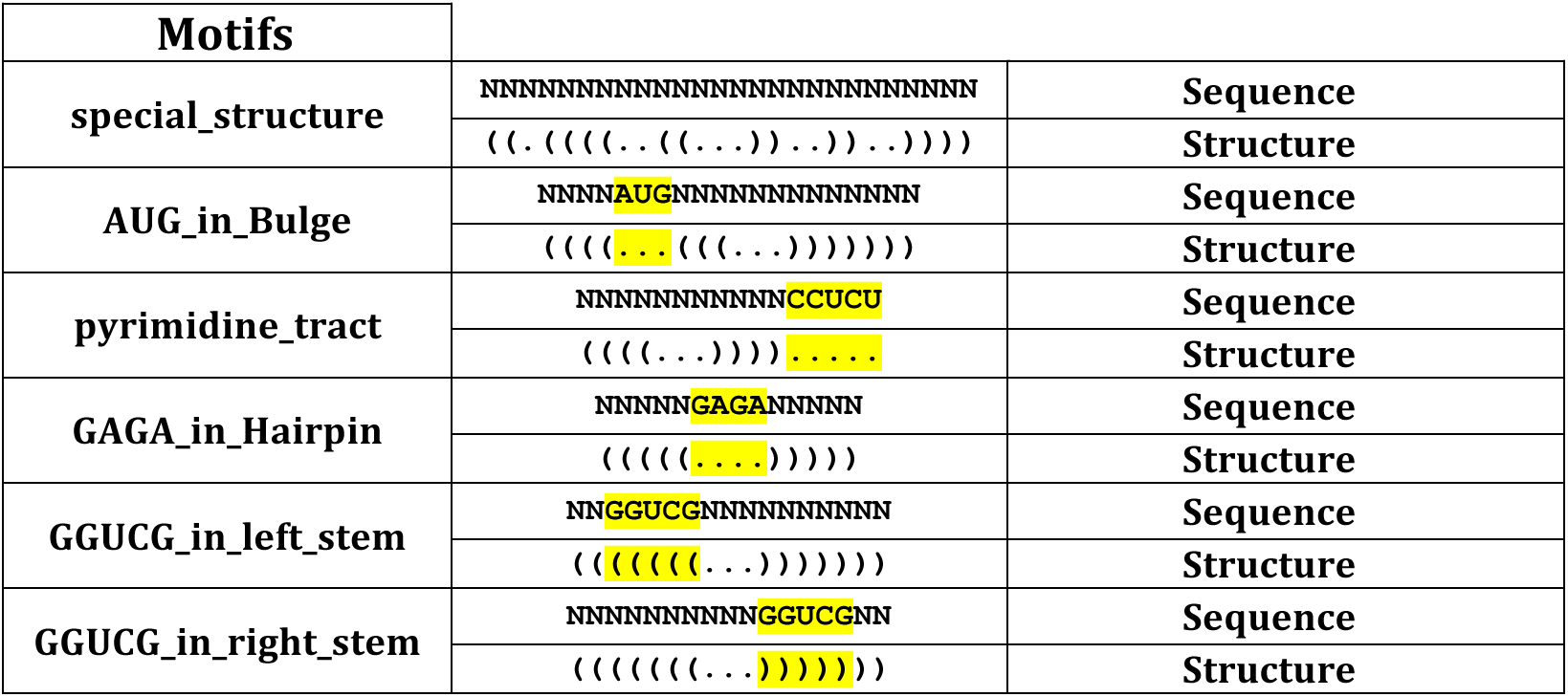
Synthetic motifs used to test SARNAclust

First, we tested RNApeakFold on synthetic CLIP peaks. These peaks were generated by adding 20 nucleotides uniformly randomly both 5’ and 3’ of the 600 synthetic sequences mentioned above. This accounts for the fact that experimental CLIP peaks contain additional nucleotides flanking the true binding motifs. RNApeakFold was used to predict secondary structures after adding +/− 100 uniformly random flanking nucleotides. Table 2 shows the comparison between using RNAfold on the simulated CLIP peaks and RNApeakFold in terms of how many of the synthetic sequences had a correct structure prediction. Note that RNAfold did not predict a single structure correctly, while RNApeakFold performed well in multiple motif types, especially *GAGA_in_Hairpin, GGUCG_in_left_stem* and *GGUCG_in_right_stem*.

**Table 2:** Comparison between RNAfold and RNApeakFold for structure prediction of synthetic CLIP peaks. Values indicate the number of structures correctly predicted by each method for each synthetic motif type.

| motifs | RNAfold | RNApeakFold | Total |
| --- | --- | --- | --- |
| **GAGA_in_Hairpin** | 0 | 77 | 100 |
| **GGUCG_in_left_stem** | 0 | 82 | 100 |
| **GGUCG_in_right_stem** | 0 | 76 | 100 |
| **pyrimidine_tract** | 0 | 12 | 100 |
| **AUG_in_Bulge** | 0 | 10 | 100 |
| **special_structure** | 0 | 0 | 100 |

Although these results are promising, we note three potential limitations of the method. First, and common to all structure prediction methods, if the real motif or the structural context is longer than the peak then the prediction will be incorrect. Second, the method may be prone to overestimating structure. That is, although it can effectively predict structured regions, it might also predict structure where single strandedness is correct. And third, it assumes that the relevant structure is stable enough to be robustly found despite the random flanking sequence. The method will not be suitable if the structure requires a specific (unknown) context around it. We observed signs that the method is affected by structural stability, as shown in the results in Table 2. The structures of the motifs that are inherently more stable (for instance *GAGA_in_Hairpin*) are predicted more accurately, while those that are inherently unstable (*special_structure*) are not.

Next, in order to test the motif identification aspects of SARNAclust, we performed clustering with all 1600 sequences. To cluster we used the EdEN graph kernel surveying over possible values for the threshold parameter, which specifies the minimal similarity for two data points to be in the same cluster. Figure 3 shows the different Fowlkes-Mallows (FM) index (Fowlkes and Mallows 1983) values for each graph transformation and each *threshold* value. The Fowlkes-Mallows index computes the similarity between the clusters returned by the clustering algorithm and the benchmark classifications, with higher FM values indicating greater similarity. It can be computed using the formula:

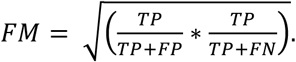

**Figure 3.**
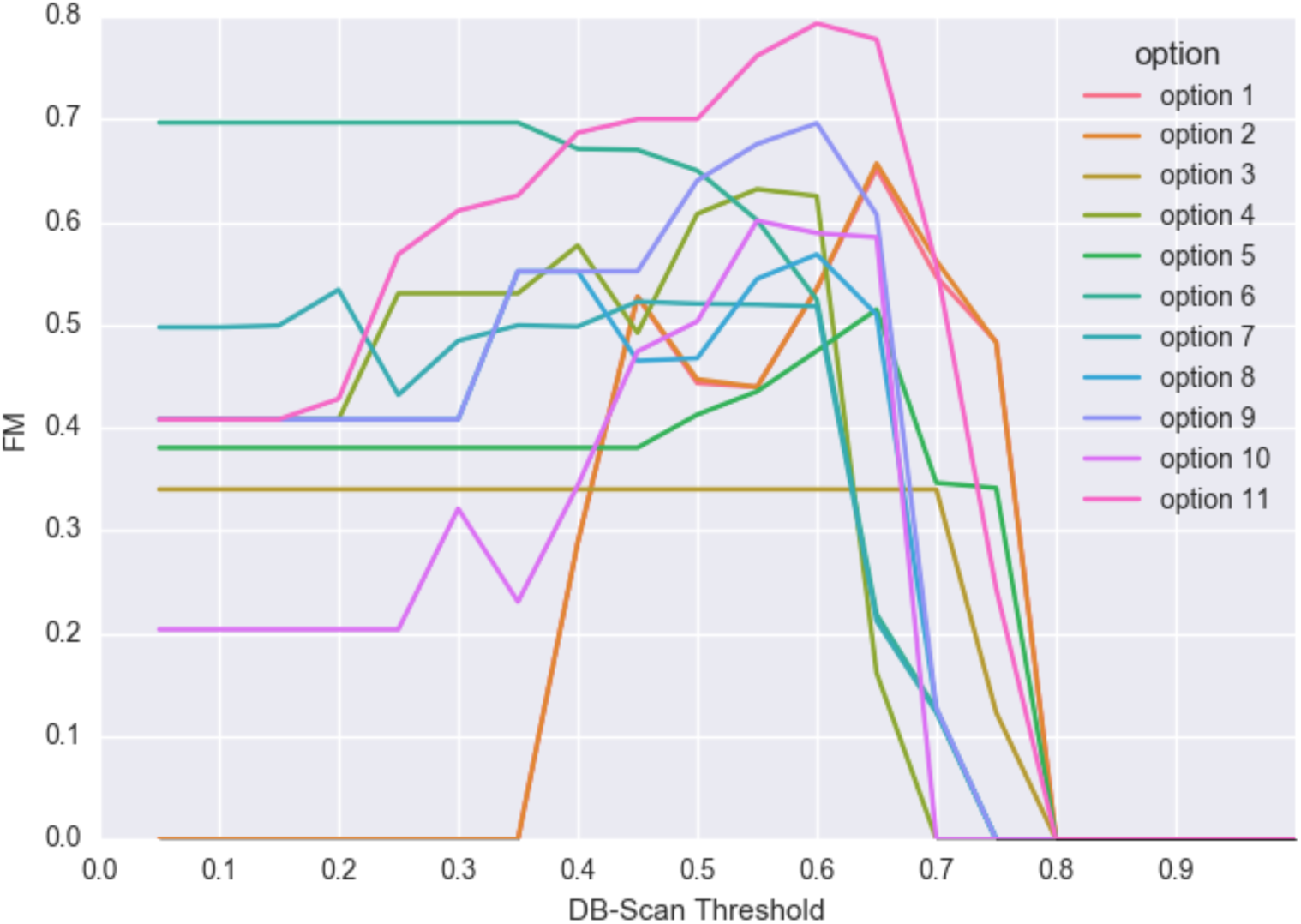
<u>Legend</u>:Fowlkes-Mallows index at different threshold values for each of the graph transformation options.

TP is True Positives (all non random sequences clustered in the same cluster; however if 2 non random motifs are clustered together then those are considered as FNs). FP is False Positives (all random sequences that appear in clusters). FN is False Negatives (all the non random sequences that do not appear in consistent clusters). True Negatives are the random sequences that do not appear in any cluster. Complete statistics for each choice of graph transformation can be found in Supplementary Data 2.

As can be seen, option 11 achieves the best FM value at *threshold* 0.6, followed by option 6 at low *thresholds* and option 9 also at *threshold* 0.6. The main conclusions of this analysis are the following (see also Supplementary Data 2):

- GraphProt-like options perform relatively well at high *threshold* values, but cannot successfully cluster at low thresholds. This is due to the excess information (i.e. number of features) specified in this graph transformation, making it difficult to find clusters of sequence/structures unless the instances are nearly identical. Option 2 contains slightly fewer features than option 1 and thus performs slightly better.
- Each graph transformation finds its corresponding motif at low to mid threshold values before suffering an increase in False Positives as threshold increases. For instance, option 4 finds the GAGA_in_hairpin motif easily at low thresholds.
- Results for option 6 are misleading since it profits from the fact that most motifs do not have external loops. So for most motif instances it will only be able to use the bulge graph features to discriminate motif instances from one another. At the same time it prevents False Positives from the random sequences that do have external loops.
- Option 3 is one of the few capable of clustering peaks of the special structure motif, but its performance is poor in general.
- Option 10 is one of the few capable of clustering peaks with the GGUCG in the stem region. It presents comparable results to the GraphProt-like options while also yielding successful clustering at low thresholds.
- Option 9 has one of the best results and is capable of finding almost all motifs with relatively low false positive rates.
- Option 11 performs the best among all options. However, this option specifies more features than option 9, and therefore it will be more affected by noise within the motifs, an issue similar to overfitting. We prognosticate that against real noisy data the performance would drop significantly.

For all these reasons, we believe that option 9 at *threshold* 0.6 is the best option moving forward. However, option 11 needs to also be considered and option 10 is appropriate if we have reason to believe there is a sequence constraint in a double-stranded region. The rest of the options can be used if the structural context of the motif is known beforehand. For instance, if we expect our motif would be found in an external loop, then we could use option 6.

### Identification of RNA motifs in ENCODE CLIP Data

In order to test our pipeline and new tools we analyzed data from ENCODE. ENCODE is conducting ongoing assays of RNA crosslinking immunoprecipitation that are expected to eventually cover > 200 known human RNA Binding Proteins using the most recent CLIP experimental protocol: eCLIP (Van Nostrand et al. 2016). We downloaded a set of 30 RBPs from ENCODE eCLIP experiments at www.encodeproject.org, each with 2 replicates and a control. For proteins with more than one experimental cell type, we used the data from the K562 female cell line. We started from the ENCODE bam files and called peaks as indicated in the methods section.

We first assessed the percentage of peaks that fall into each type of genomic region: Coding Region, 5’UTR, 3’UTR, intronic region or non-coding RNA. Figure 4 shows the region distribution for the union of CLIP sites for all RBPs (Fig. 4A), as well as the distribution values averaged across RBPs (Fig. 4B). These two measures yield similar results, and both show that most peaks fall into intronic regions. Potential explanations are that most proteins are involved in splicing, RNA structure or localization; that RBPs bind promiscuously; and/or that eCLIP is noisy and that such peaks are due to non-specific signal from intronic RNA.

**Figure 4.**
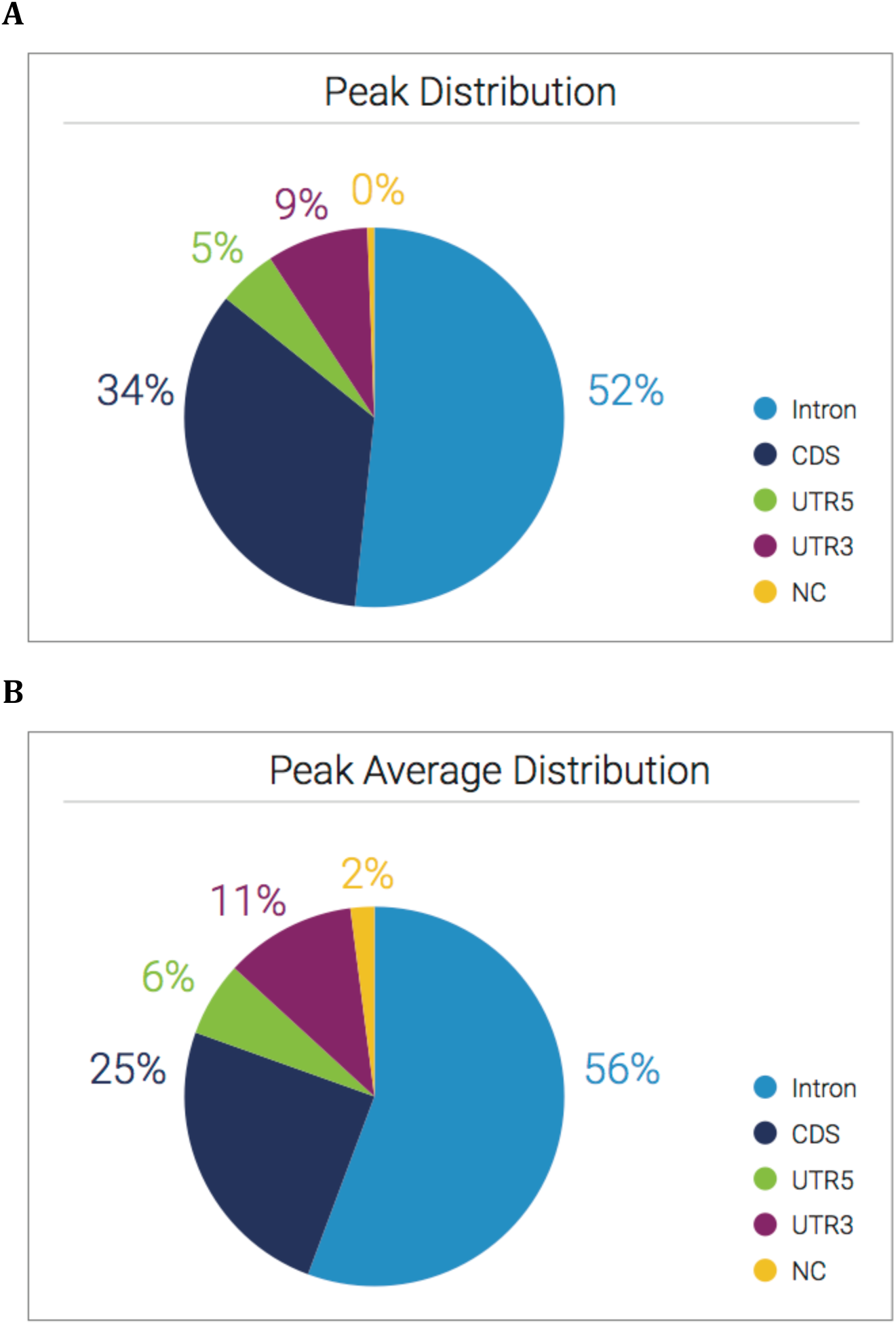
<u>Legend</u>: Pie chart of ENCODE eCLIP peaks composition. Regions are divided into intronic, non-coding, 5’ UTR, CDS and 3’UTR. In a) peaks from all RBPs are pooled together in order to calculate the global composition. In b) compositions are calculated for each RBP, and the pie chart shows the average across RBPs.

As an initial analysis of motifs in the ENCODE data, we searched for common k-mers in the peaks in each set. This approach ignores RNA structure, providing a comparison to the sequence/structure clustering approach. We performed k-mer analysis as in (Yeo et al. 2009, 2) for k = 4,5,6,7,8,9. Significant k-mers (Z-score > 2.5) are shown in Supplementary Data 3. For comparison, we applied SARNAclust but without incorporating structural information. For each of the ENCODE RBPs, we calculated the best motif from the k-mer analysis and from this SARNAclust structureless analysis (Table 3). Motifs that have been previously reported for these proteins as found in RBPDB and ATtRACT (Giudice et al. 2016), as well as the RNA binding domain type, are also shown in Table 3 for comparison. For each protein, the best k-mer was chosen as the one that maximizes the quantity (Z-score)*k.

**Table 3:**
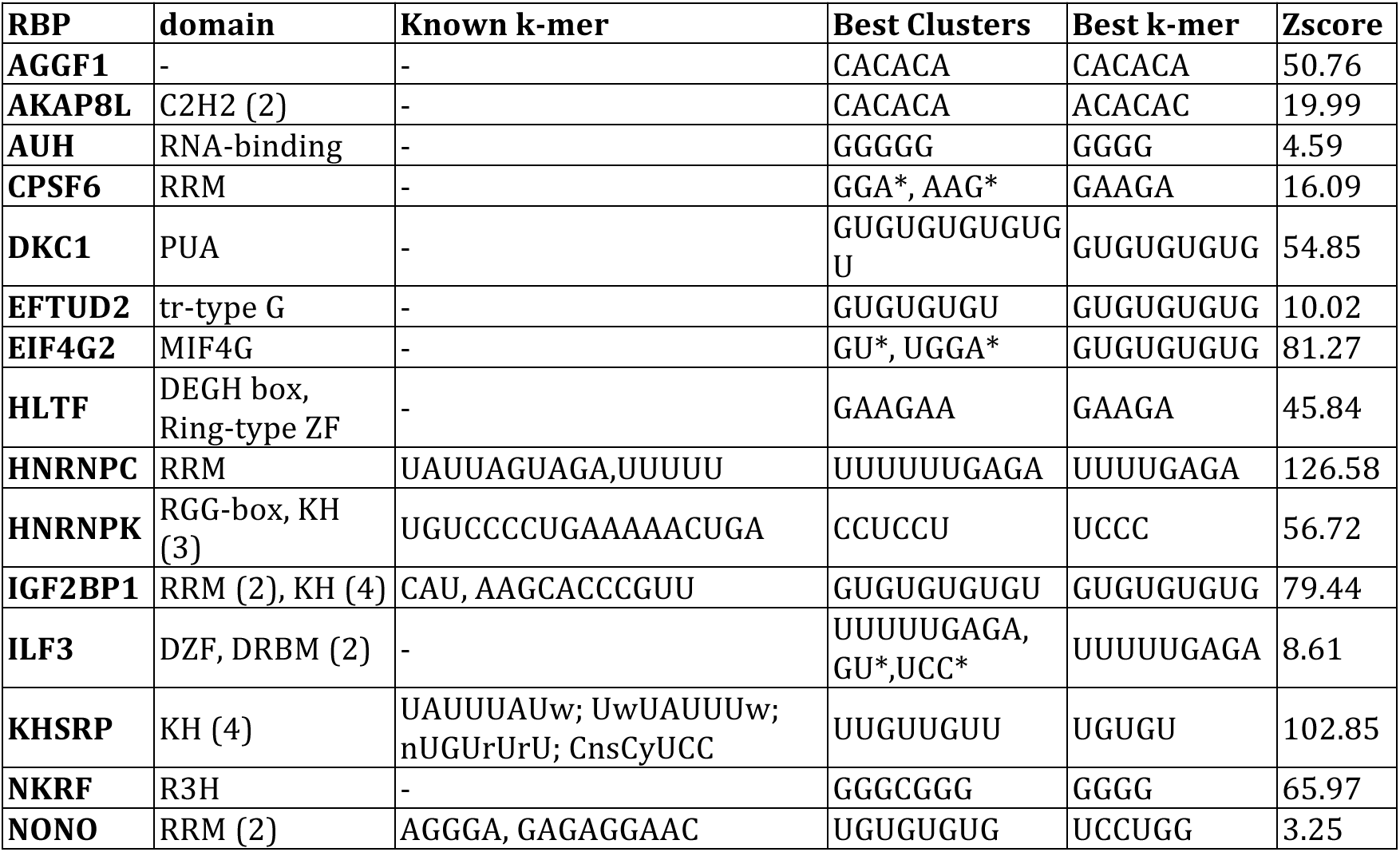
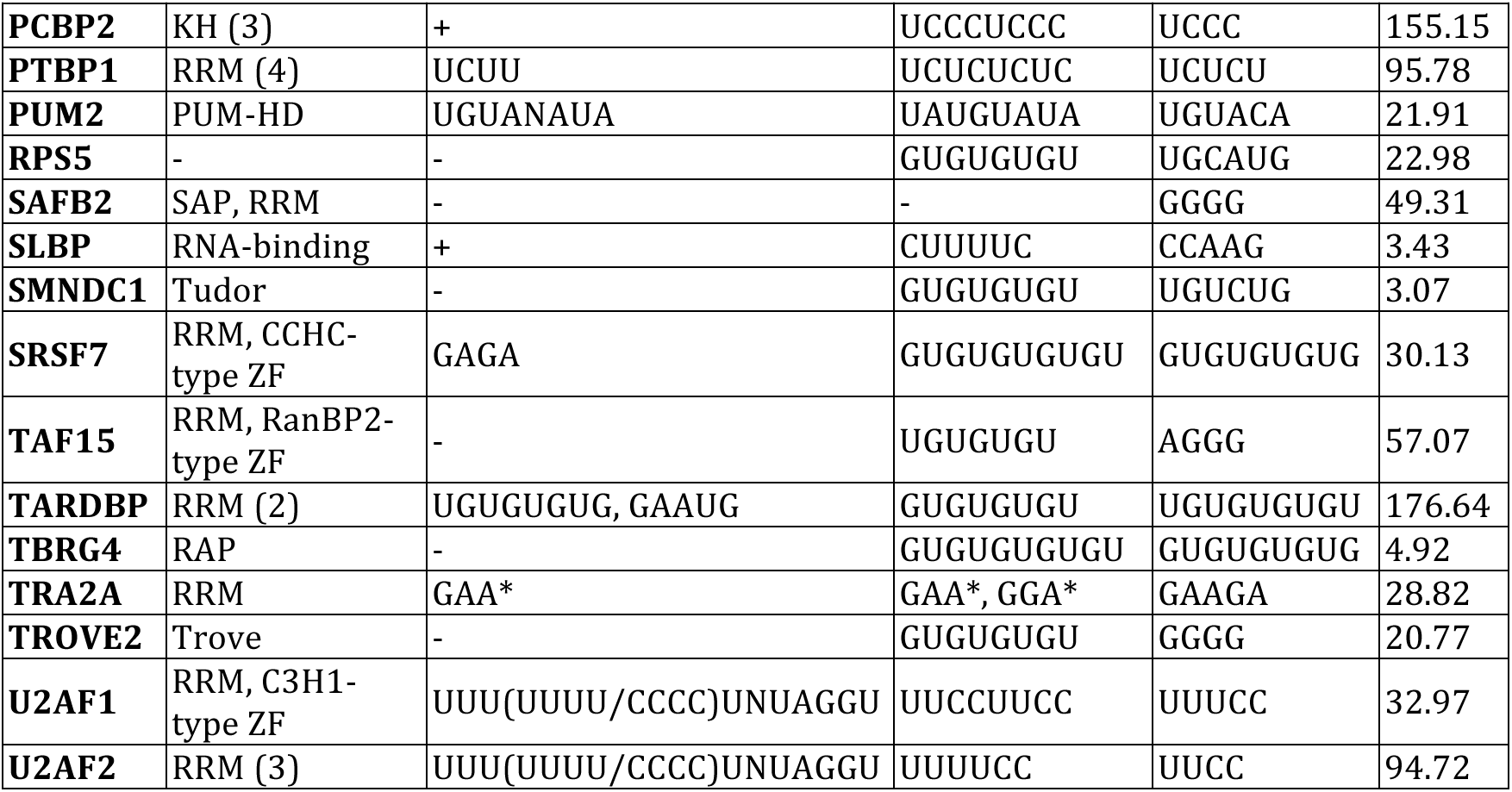
Known and predicted motifs for selected RNA binding proteins. For each RBP, the known binding domain, literature known k-mer, SARNAclust prediction, k-mer prediction, and k-mer z-score are shown.

There are several conclusions we can extract from this table:

- Most of the RBPs selected for this study have motif predictions dominated by low complexity k-mers. Many are G-rich, GU repeats, GGA repeats, or GAA repeats, possibly indicating experimental noise or promiscuous binding. For the RBPs with a significant k-mer, the k-mer with the best score is often a subsequence of a longer statistically significant k-mer with a worse score.
- In most of the cases, SARNAclust finds motifs similar to the best k-mer. However, SARNAclust yields these without requiring one to survey over values of k.
- Some RBPs show binding to CU rich regions, matching prior literature descriptions (Song et al. 2012; Toyoda et al. 2007; Yuan et al. 2002).
- Four RBPs show binding to CA repeats, even though they do not share the same RNA binding domains.
- In general, there is no correlation between RNA binding domain class and the number of significant k-mers.
- Several proteins yield specific motif predictions, previously known or otherwise, such as ILF3, HNRNPC, SLBP and PUM2.

We also found that recalculating clusters after removing intronic peaks did not alter the results significantly. Therefore intronic peaks are not unusual and they do not, in general, dominate k-mer or cluster analysis.

Next, we analyzed the effect of adding RNA structure to the motif identification process. To do this we used RNApeakFold to calculate the secondary structure around each peak and applied SARNAclust with graph transformations 9, 10 or 11 at similarity *threshold* 0.6, based on our calibrations from the synthetic data. However, we expected that for real data there would be more noise and false positives. Noisy data in general cause SARNAclust to yield more low information content clusters, i.e. clusters that cannot be coherently grouped by sequence or structure.

Table 4 shows SARNAclust results for all the ENCODE RBPs which yielded significant clusters. RBPs not in this table either yielded no clusters or only low information content clusters. Most RBPs show motifs similar to either their known motifs or the ones found in the pure sequence analysis. Also, as expected, option 11 was less effective at finding significant clusters, and option 9 was the clear winner. Interestingly, option 10 yielded a strong prediction for the SLBP motif, coinciding with the literature-known motif (Battle and Doudna 2001). This prediction indicated that sequence conservation within the base paired region of the RNA site is important for SLBP protein binding. Another interesting observation was the identification of a motif for ILF3. This motif was similar to that found when no structural information was used, and it was also similar to the known motif for splicing involved RBP HNRNPC (Zarnack et al. 2013). In addition, table 4 indicates a variety of other results, such as the frequent importance of GU* (GU repeats) for several RBPs, novel motifs for CSPF6 and SAFB2, and alternate predicted motifs for TRA2A.

**Table 4:**
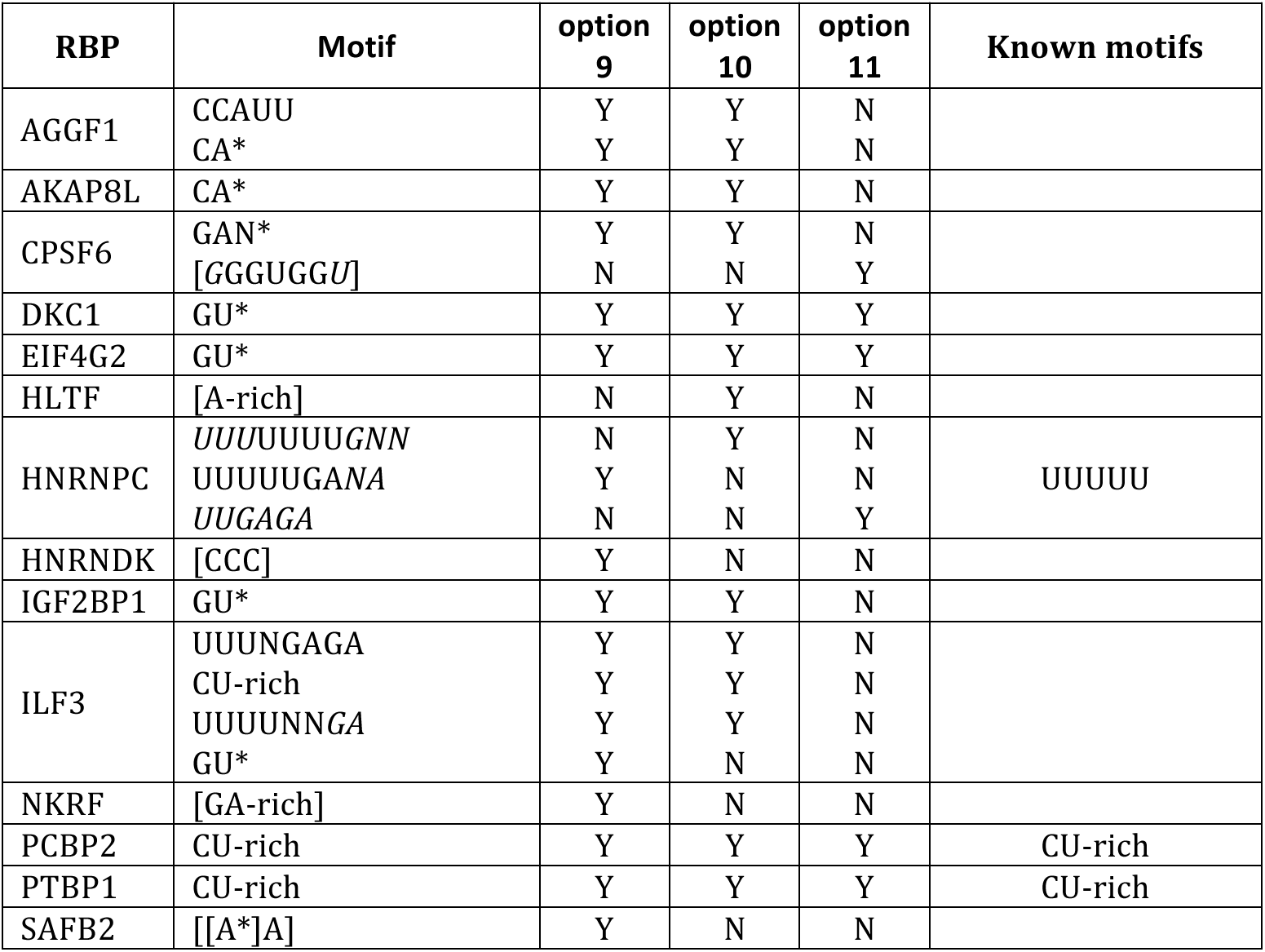
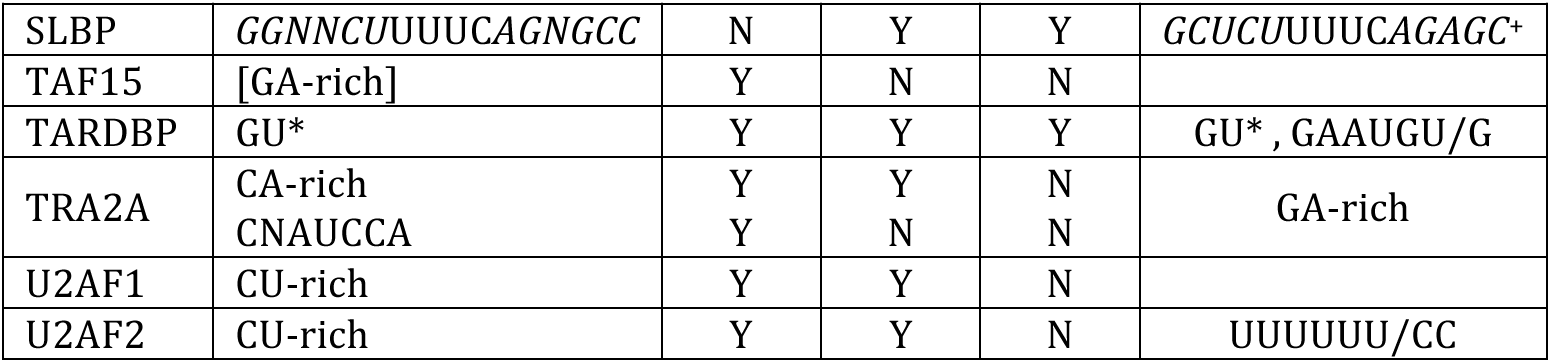
SARNAclust results for RBPs that yielded significant clusters. Y/N indicate whether the motif was detected with each option choice. Nucleotides in *italic* indicate they are base paired. [indicates start or end of a stem. + from (Battle and Doudna 2001)

### Experimental validation of SARNAclust prediction for SLBP

As described above, SARNAclust yielded a clear prediction for the RNA motif of SLBP, matching the literature-annotated version of the motif (Battle and Doudna 2001). Moreover, part of the signal for this motif arose from sequence conservation in the double stranded region. To determine if SARNAclust could effectively predict the importance of features within motifs, we performed additional experiments of SLBP-RNA binding using designed sequences in a RNA bind-n-seq (RBNS) validation protocol. To do this, we first used RNAiFold (Garcia-Martin et al. 2015) (See Methods) to design four different types of sequences as illustrated in Table 5. These sequences were chosen to determine whether sequence conservation is necessary in both the loop region and the stem region of the motif, and also if the occurrence of sequence of the loop region in a different structural context, i.e., in a bulge, could still lead to protein-RNA binding. We then performed RBNS using purified GST-SBP-SLBP to pull down the designed RNA sequences (Lambert et al. 2014). As a nonspecific binding control, we also performed RBNS with the same RNA against purified GST-SBP. RBNS was performed in duplicate for each protein tested. Each of these proteins were expressed in *E. coli* and affinity purified to sufficient purity (Supplementary Figure 3). RNA pulled down by each protein was reverse transcribed with a primer containing a 10 nucleotide random sequence to collapse PCR duplicates during data analysis. The resulting cDNA was then PCR amplified to attach Illumina sequencing primers and indices.

**Table 5:**
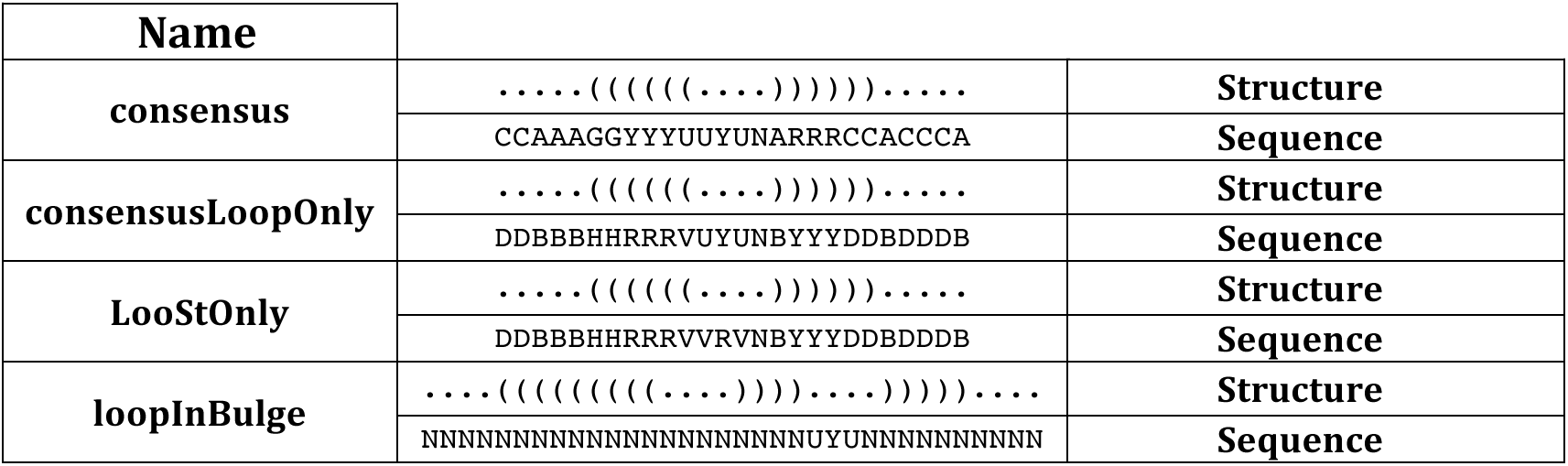
Four different classes of designed sequences used for the RNA Bind-N-Seq validations for SLBP.

Figure 5 shows the shift in percentage of reads of each type and its difference between GST-SBP RBNS and GST-SBP-SLBP RBNS. Only the consensus motif from (Battle and Doudna 2001) has a clear shift from the control, indicating that the motif definition is specific and that the variant versions of the motif have decreased binding. To assess p-values, we used DEseq (Anders and Huber 2010) to compare the readcounts (Supplementary Data 4) of all sequences in the pool to the controls. Here an increased normalized ratio of counts in the SLBP pulldown relative to the control indicates increased binding. This analysis showed that only sequences from the consensus motif bind to SLBP significantly. Moreover, all but 7 of these consensus sequences are significantly overrepresented in the SLBP bound pool (Adjusted p-val > 0.01). These results indicate that disruption of any of the elements in the SARNAclust prediction for the SLBP motif decrease binding. Furthermore, Supplementary Figure 4 shows the sequence logos for all the consensus sequences that bind or do not bind significantly, respectively. The logos indicate that long stretches of U’s near the apical region of the hairpin loop compromise binding affinity, which is to be expected since they are energetically unfavorable and therefore prone to render the hairpin unstable.

**Figure 5.**
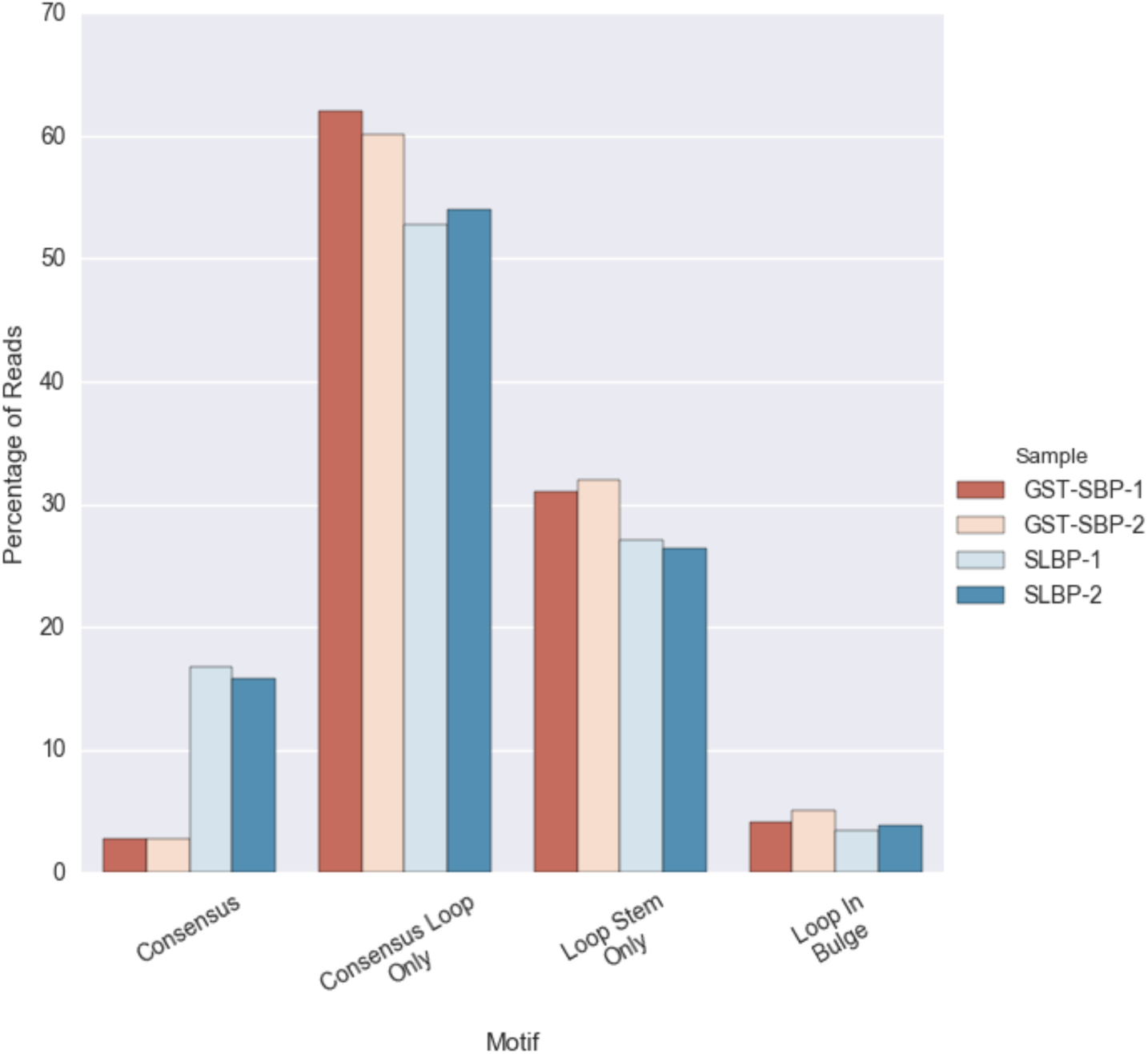
<u>Legend</u>: Percentage shift in the sequences retrieved in samples versus control for SLBP sequences.

To further validate these results we performed several gel shift experiments (Figure 6). We incubated 6 RNA probes selected from the RBNS data with purified GST-SBP-SLBP. Supplementary Data 4 shows the 6 selected sequences highlighted in red. These include 2 from the consensus binding group, one with strong binding affinity in the RBNS assay (consensus A) and one with no significant binding affinity (consensus B). There are also 4 extra sequences from the remaining types where the RBNS binding signal was not significant. As expected, only the consensus A sequence shows binding to SLBP, confirming our conclusions from the p-value analysis and validating the RBNS protocol.

**Figure 6.**
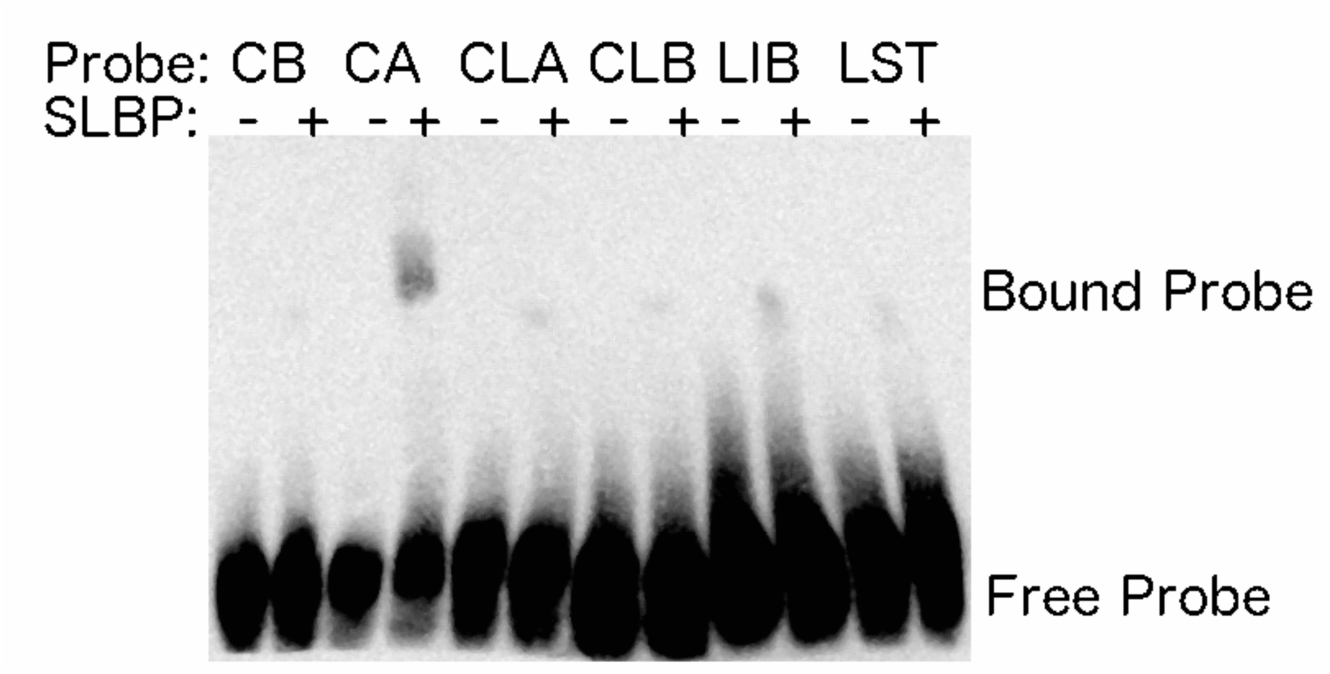
(gel shift for SLBP) <u>Legend</u>: Gel shift results for select probes tested in the RBNS when incubated with purified GST-SBP-SLBP. The Consensus A (CA) probe shows more binding relative to Consensus B (CB), Consensus Loop Only A (CLA), Consensus Loop Only B (CLB), Loop In Bulge (LIB) and Loop Stem Only (LST). Sequences for each probe and their RBNS results can be found in Supplemental Data 4.

### Identification of a Novel Motif for ILF3

We next used the RBNS approach to test a newly predicted motif for the protein ILF3, which was predicted by SARNAclust to have a motif similar to one previously described for the protein HNRNPC (Zarnack et al. 2013). In particular, we used RBNS to test the binding of the predicted motifs with ILF3 and whether it requires a specific RNA structure.

Using RNAiFold we generated sequences for 4 different perturbations of the motif as shown in Table 6. Similarly to SLBP, we performed RBNS with purified GST-SBP-ILF3 using an RNA pool based on the motifs in Table 6 (Supplementary Figure 3). Figure 7 shows the shift in percentage of reads of each type and its difference between GST-SBP-1, GST-SBP-2 controls and ILF3-1, ILF3-2 samples. It can be seen that only the motif *uuuugaga-paired* has a shift from the control. This indicates that the novel motif is real. We also used DEseq to analyze differential representation of sequences in the ILF3 bound and unbound pools (Supplementary Data 5). Only sequences from the *uuuugaga-paired* motif bind to ILF3 significantly, confirming and specifying the computationally discovered motif. These results indicate that ILF3 binds to an UUUUGAGA motif with most nucleotides in double-stranded regions. This supports a relationship between ILF3 and HNRNPC, which has been reported to have a very similar motif. Our experimental finding that the motif is likely to occur in a double stranded region is novel, as this motif has not been previously reported for ILF3. We note that the SARNAclust result for ILF3 showed some indications of double strandedness, but not decisively so. It is worth mentioning that SARNAclust also predicted a separate GU repeat motif for ILF3 (*gu-repeats* motif in Table 6), but the motif did not show enriched binding. GU-rich motifs were predicted for many other ENCODE RBPs as well, and we speculate that the presence of such sequences in CLIP data may be due to experimental noise.

**Table 6:**
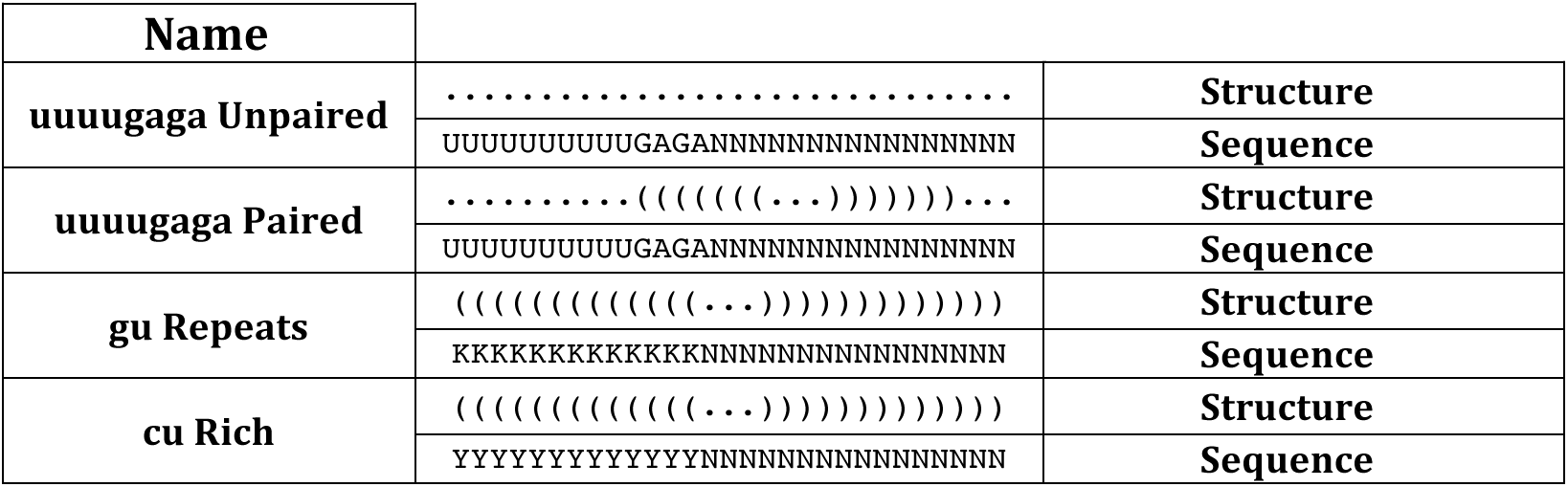
Four different classes of designed sequences used for the RNA Bind-N-Seq validations for ILF3.

**Figure 7.**
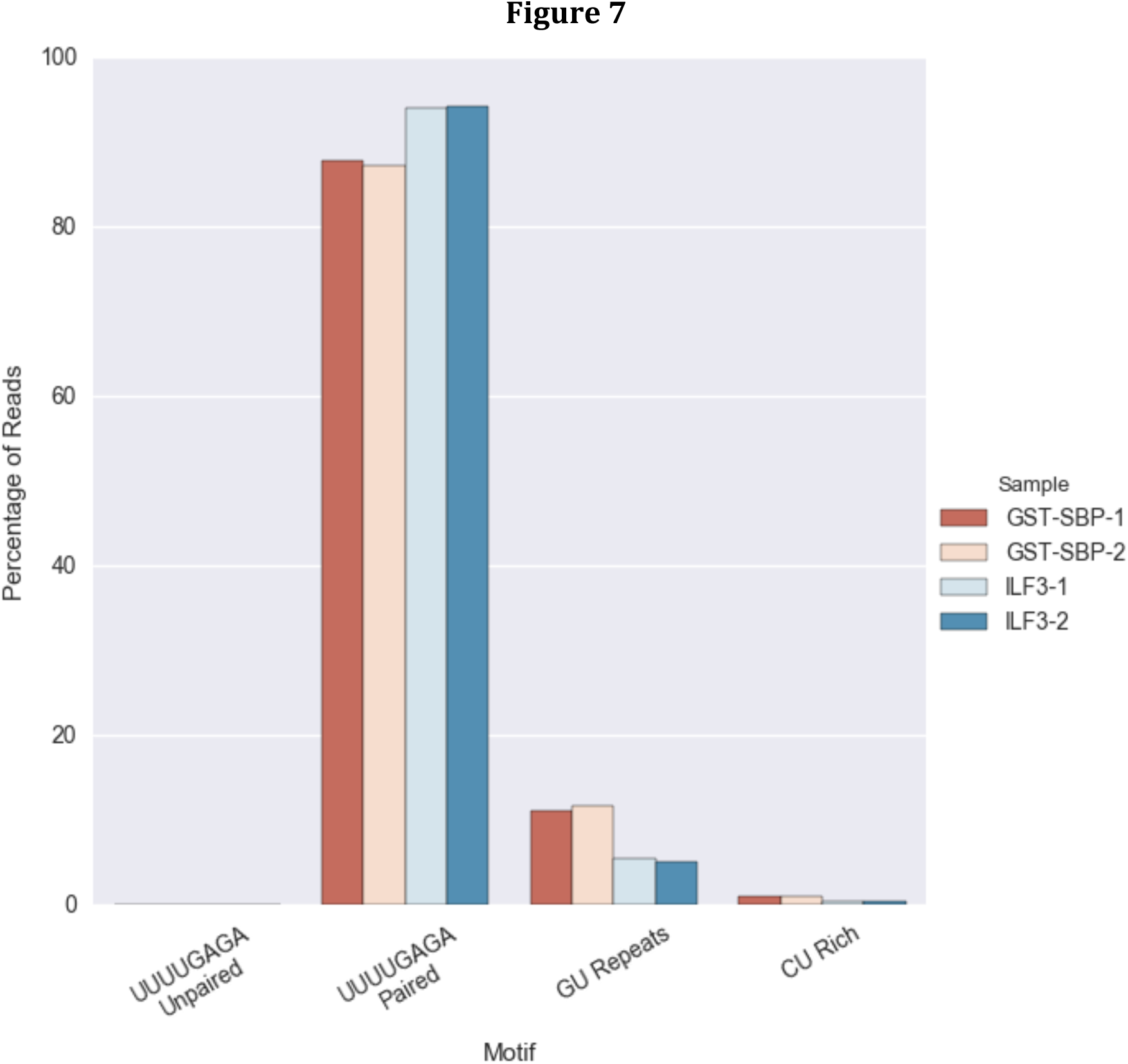
<u>Legend</u>: Percentage shift in the sequences retrieved in samples versus control for ILF3 sequences.

## Discussion

SARNAclust is a novel computational method that can effectively process and analyze data from CLIP experiments in order to predict RNA motifs likely to bind individual proteins. A key novelty of SARNAclust is that it can assess RNA binding motifs at the level of the complete RNA structure, rather than only taking into account abstractions of structural context. SARNAclust makes use of RNApeakFold, which we have developed and tuned to estimate structure at CLIP sites. This is a key component of the pipeline, as other structure prediction approaches such as RNAfold do not correctly recapitulate the structures in our synthetic data. Even without using structure information, we have shown that SARNAclust is able to identify motifs as effectively as k-mer analysis, and when structural information is added SARNAclust is able to further identify known motifs and previously unknown motifs. The SARNAclust approach of clustering rather than classifying distinguishes it from prior methods, allowing it to identify motifs even without training data. This is an important aspect for CLIP-seq, for which the specificity of experimental measurements is not well understood due to diverse effects such as multiple binding modalities and sources of noise.

From the analysis of the ENCODE data with SARNAclust and RNA Bind-n-Seq, we can make several observations about RNA-protein binding. First, although a great number of CLIP peaks fall in intronic regions, motif predictions are not substantially biased by them. This suggests many intronic regions contain real motif instances. Second, there are a great number of G-rich and GU repeats in the motifs of different RBPs, suggesting these may be non-specific signals. For ILF3 we were able to experimentally verify binding to a newly predicted UUUUGAGA motif, but not to repetitive GU motifs, supporting the idea that those repetitive sequences are not true binding sequences. Third, we observed no relationship between protein binding domain and motif, indicating substantial protein-specificity in binding behavior. Fourth, SARNAclust allowed us to investigate the relative importance of structure, which has been challenging for RNA-protein interactions, and we found that structure significantly affected the Bind-N-Seq results for both SLBP or ILF3. Structural changes to each of several components of the SLBP motif reduced binding, and the new motif for ILF3 exhibited a bias for double-strandedness.

The similarity of the new ILF3 motif to that for HNRNPC is intriguing, as it was shown in (Zarnack et al. 2013) that HNRNPC competes with another protein U2AF2 for binding of 3’ splice sites to regulate the inclusion/exclusion of exons. They concluded that HNRNPC prevents inclusion of cryptic exons while U2AF2 promotes it, with RBP binding often occurring in antisense Alu elements. Based on this competition, we would expect U2AF2 to have a similar binding site to HNRNPC. However, the predicted motif for HNRNPC is much more similar to that for ILF3 than it is to the predicted U2AF2 motif (Tables 3 and 4). This suggests that ILF3 might compete with either HNRNPC or U2AF2 for binding of similar regions. Supplementary Figure 5 shows the overlap of binding sites between ILF3 and both HNRNPC and U2AF2. The overlap with HNRNPC is greater than that of U2AF2. Moreover, if we restrict the sites to those that overlap with anti-sense Alu elements, this difference is amplified (Supplementary Figure 6). This might indicate that ILF3 is competing with HNRNPC for binding in anti-sense Alu regions. This hypothesis is further supported by the large overlap of both HNRNPC peaks and ILF3 peaks with anti-sense Alu regions, while there is less overlap of U2AF2 peaks with the anti-sense Alu regions (Supplementary Figure 7). Further experimental studies will be needed to ascertain this hypothesis.

In summary, we have introduced a new pipeline SARNAclust for analyzing CLIP data in order to cluster CLIP peaks into different binding motifs. We have also extensively analyzed 30 recent eCLIP experiments performed by ENCODE, both with traditional k-mer analysis and using our new algorithms. We have verified the effectiveness of SARNAclust on synthetic data and used RNA Bind-n-Seq to experimentally validate predictions for new and known binding motif predictions in the ENCODE data. In the future, and as more eCLIP data sets for double-stranded binding RBPs become available, we expect SARNAclust will be a valuable tool to discover new motifs, probe the combinatorial interactions of RNA-binding proteins, and elucidate their functional importance. SARNAclust is available on Github at https://github.com/idotu/SARNAclust.

## Methods

### a. ENCODE data

ENCODE data (www.encodeproject.org) correspond to a set of CLIP experiments described as enhanced CLIP (eCLIP), which modifies the iCLIP method to include improvements in library preparation of RNA fragments. See (Van Nostrand et al. 2016) for details. All data were downloaded through the ENCODE Project website.

### b. Bam to peaks file processing

We calculated the set of clusters or peaks for each RBP by running pyicoclip on the ENCODE bam files (2 replicates each). The software pyicoclip is part of the pyicoteo software for analysis of high-throughput sequencing data (Althammer et al. 2011) (available at https://bitbucket.org/regulatorygenomicsupf/pyicoteo). Pyicoclip implements the modified False Discovery Rate approach proposed in (Yeo et al. 2009, 2) to determine significant clusters in a list of genomic regions. Pyicoclip implementation, together with the pyicoteo software, offers a flexible and effective framework for the processing and analysis of different types of CLIP-Seq data, with or without associated controls. We chose pyicoteo for its speed and because its modular architecture allowed us to adapt the CLIP-Seq analyses for data standardization. In order to generate a final set of peaks for each RBP, we used peaks that overlapped both replicates and subtracted peaks overlapping with the control. We chose this approach rather than using enrichment thresholds in order to minimize noise from any systematic measurement biases. For each peak we tracked the gene it overlapped, the type of region within the gene, and the genomic sequence in the +/− 100 flanking nucleotides.

### c. Structure Prediction

Given an RNA sequence S (of length n) of a peak along with +/− 100 flanking nucleotides, we calculate the structure of the peak following Nussinov recursions:

> Init (for i in 1 to n)
>
> 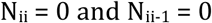
>
> Recursion (for i,j in 1 to n)
>
> 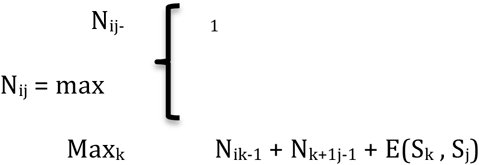

E(S_k_, S_j_) is the base pairing probability as calculated using RNAfold –p from the Vienna package [31] on the complete sequence (peak +/− 100 flanking nucleotides),

### d. Clustering algorithm

The clustering algorithm is the main component in our pipeline. Given a set of RNA sequences along with their predicted secondary structures, it identifies clusters of similar RNAs by encoding both sequence and structure as a graph, and using the EDeN kernel similarly as in GraphClust. The pipeline accepts several parameters to control both the graph transformation and the clustering. It also allows for the use of only sequence information in a sliding window fashion. The clustering algorithms supported are: K-means, Mean Shift, DB-Scan, Affinity Propagation and Spectral Clustering from sklearn package (http://scikit-learn.org/stable) and Density Clustering (Rodriguez and Laio 2014) in an in-house implementation. A full implementation is available at the Github site. For the clustering of the synthetic motif data we used the EdEN graph kernel with DB-SCAN, surveying over possible values for the DB-Scan parameter *threshold* (http://scikit-learn.org/stable/modules/generated/sklearn.cluster.DBSCAN.html), which specifies the minimal similarity for two data points to be in the same cluster. Other parameter choices were *radius* = 2 and *distance* = 2 with *min_samples* = 10 (see (Heyne et al. 2012)),

### e. RNAiFold to generate candidates

In order to generate candidate RNA sequences for our RBNS experimental validation we used the RNA inverse folding software RNAiFold (Garcia-Martin et al. 2015). Given a sequence/structure RNA motif, we generated thousands of sequences that fold into the given secondary structure and maintain the given sequence constraints. Sequences generated by RNAiFold were used in the design of the RBNS pool. Moreover, we used RNAiFold to generate sequences corresponding to perturbations of the SLBP and ILF3 motifs. This was done by altering constraints and re-running RNAiFold, e.g. for SLBP we moved a sequence motif from a stem loop to a bulge to generate a pool of sequences that would test whether location of the sequence motif within the structure affected binding.

### f. Experimental Protocols

#### *In vitro* protein expression and purification

A previously generated pGEX6P1-based expression vector containing streptavidin binding peptide (SBP)-tagged ILF3 was used for ILF3 binding experiments. For SLBP experiments, the protein coding sequence for SLBP was codon optimized for *E. coli* expression using the IDT Codon Optimization Tool and Gibson assembled into pGEX6P1-SBP, which enhanced solubility compared to the human sequence. Each plasmid was transformed into Rosetta(DE3) pLysS *E. coli*. Protein expression was induced with 1mM isopropylthiogalactoside (IPTG) and grown for 4 hours at 16°C. Soluble protein was extracted from the bacteria using the Qproteome Bacterial Protein Prep Kit (QIAGEN). The proteins were then affinity purified using Glutathione Sepharose 4B and eluted in a buffer containing 0.2% Triton X-100 and concentrated using Corning^®^ Spin-X^®^ UF with a 10 kDa molecular weight cutoff (MWCO). Proteins were then equilibrated into RBNS binding buffer (25mM Trish ph 7.5, 150mM KCl, 0.1% Tween, 0.5 mg/mL BSA, 3mM MgCl_2_, 1mM DTT) using Zeba desalting columns 7KDa MWCO. Purified proteins were then frozen at -80°C for short-term storage. Protein concentrations were obtained using Pierce^™^ BCA Protein Assay Kit. Protein purity was assessed using SDS-PAGE.

#### RNA pool generation for RNA Bind-N-Seq

Oligonucleotide sequences were ordered from CustomArray Inc. in a 12,472 oligo pool. PCR was used to amplify the ILF3 pool (5’- CCCATAATACTTGTCCCG -3’ and 5’- TAATACGACTCACTATAGGG-3’) and the SLBP pool (5’- CTTGACTGCGAGCTGTTGA-3’ and 5’- TAATACGACTCACTATAGGTCACGTC-3’). *In vitro* transcription of the oligo pool was performed using a AmpliScribe^™^ T7 High Yield Transcription Kit. The RNA was purified by lithium chloride precipitation and resuspended in RBNS binding buffer.

#### RNA Bind-N-Seq

RBNS was performed as described in (Lambert et al. 2014). 27 nM of each protein was incubated with 750 pM of RNA. RNA was reverse transcribed using SuperScript^®^ III Reverse Transcriptase and a primer containing a 10 nucleotide barcode for SLBP (5’- GTGACTGGAGTTCAGACGTGTGCTCTTCCGATCTNNNNNNNNNNCTTGACTGCGTGCTGTTGA-3’)and ILF3 (5’-GTGACTGGAGTTCAGACGTGTGCTCTTCCGATCTNNNNNNNNNNCCCATAATACTTGTCCCG-3’). PCR was performed to amplify cDNA derived from the RBNS RNAs and attach Illumina flow cell binding sequences and indices (5’- AATGATACGGCGACCACCGAGATCTACAC-i5_index-ACACTCTTTCCCTACACGACGCTCTTCCGATCT-3’ and 5’-CAAGCAGAAGACG G CATACGAGAT-i7_index-GTGACTGGAGTTCAGACGTGTGCTCTTCCGATCT-3’). DNA was sequenced on a Miseq using a 200 cycle paired end kit.

#### RBNS data analysis

FLASH was used to join paired end reads which intersected, which is expected for each of the sequences tested (Magoc and Salzberg 2011). Reads that contained the anticipated primer sequences were aligned using HISAT2 (Kim et al. 2015). Reads aligning to the same sequence and containing the same 10 nucleotide random sequence were collapsed into one read using a custom python script. The resulting counts were input to DEseq for analysis (Anders and Huber 2010).

### g. Gel Shift Experiments

Probes were *in vitro* transcribed and biotinylated using the Pierce RNA 3’ end biotinylation kit. 1 nM of biotinylated probe was incubated with either 320 nM GST-SBP-SLBP in binding buffer consisting of 10 mM HEPES (pH 7.3), 20 mM KCl, 1 mM MgCl_2_, 20 mM DTT, 5% glycerol. The incubation period was 30 minutes, followed by gel electrophoresis on a native TBE 4% polyacrylamide gel and transfer to a nylon membrane, all at 4°C.

## Acknowledgements

JHC was supported by NIH grants R21 HG007554 and R01 NS094637. EMR and EE were supported by the MINECO and FEDER (BIO2014-52566-R), AGAUR (SGR2014-1121), and the Sandra Ibarra Foundation for Cancer (FSI2013). The authors thank Gene Yeo for assistance with the ENCODE CLIP-seq data, Chris Burge and Daniel Dominguez for sharing constructs and experimental advice, and Brent Graveley for discussions.

## References

Alipanahi B, Delong A, Weirauch MT, Frey BJ. 2015. Predicting the sequence specificities of DNA- and RNA-binding proteins by deep learning. Nat Biotechnol 33: 831–838.

Althammer S, Gonzalez-Vallinas J, Ballare C, Beato M, Eyras E. 2011. Pyicos: a versatile toolkit for the analysis of high-throughput sequencing data. Bioinforma Oxf Engl 27: 3333–3340.

Anders S, Huber W. 2010. Differential expression analysis for sequence count data. Genome Biol 11: R106.

Bahrami-Samani E, Penalva LOF, Smith AD, Uren PJ. 2015. Leveraging cross-link modification events in CLIP-seq for motif discovery. Nucleic Acids Res 43: 95–103.

Bailey TL, Williams N, Misleh C, Li WW. 2006. MEME: discovering and analyzing DNA and protein sequence motifs. Nucleic Acids Res 34: W369–373.

Battle DJ, Doudna JA. 2001. The stem-loop binding protein forms a highly stable and specific complex with the 3’ stem-loop of histone mRNAs. RNA N Y N 7: 123132.

Chi SW, Zang JB, Mele A, Darnell RB. 2009. Argonaute HITS-CLIP decodes microRNA-mRNA interaction maps. Nature 460: 479–486.

Cook KB, Kazan H, Zuberi K, Morris Q, Hughes TR. 2011. RBPDB: a database of RNA-binding specificities. Nucleic Acids Res 39: D301–308.

Costa F, De Grave K. 2010. Fast neighborhood subgraph pairwise distance kernel. In Proceedings of the 26th International Conference on Machine Learning, pp. 255–262, Omnipress.

Dao P, Hoinka J, Takahashi M, Zhou J, Ho M, Wang Y, Costa F, Rossi JJ, Backofen R, Burnett J, et al. 2016. AptaTRACE Elucidates RNA Sequence-Structure Motifs from Selection Trends in. Cell Syst 3: 62–70.

Fowlkes EB, Mallows CL. 1983. A method for comparing two hierarchical clusterings. J Am Stat Assoc 78: 553–569.

Fukunaga T, Ozaki H, Terai G, Asai K, Iwasaki W, Kiryu H. 2014. CapR: revealing structural specificities of RNA-binding protein target recognition using CLIP-seq data. Genome Biol 15: R16.

Garcia-Martin JA, Bayegan AH, Dotu I, Clote P. 2016. RNAdualPF: software to compute the dual partition function with sample applications in molecular evolution theory. BMC Bioinformatics 17: 424.

Garcia-Martin JA, Clote P, Dotu I. 2013. RNAiFOLD: a constraint programming algorithm for RNA inverse folding and molecular design. J Bioinform Comput Biol 11: 1350001.

Garcia-Martin JA, Dotu I, Clote P. 2015. RNAiFold 2.0: a web server and software to design custom and Rfam-based RNA molecules. Nucleic Acids Res 43: W513–521.

Georgiev S, Boyle AP, Jayasurya K, Ding X, Mukherjee S, Ohler U. 2010. Evidence-ranked motif identification. Genome Biol 11: R19.

Giegerich R, Voss B, Rehmsmeier M. 2004. Abstract shapes of RNA. Nucleic Acids Res 32: 4843–4851.

Giudice G, Sanchez-Cabo F, Torroja C, Lara-Pezzi E. 2016. ATtRACT-a database of RNA-binding proteins and associated motifs. Database J Biol Databases Curation 2016.

Hafner M, Landthaler M, Burger L, Khorshid M, Hausser J, Berninger P, Rothballer A, Ascano M, Jungkamp A-C, Munschauer M, et al. 2010. PAR-CliP‐‐a method to identify transcriptome-wide the binding sites of RNA binding proteins. United States.

Heyne S, Costa F, Rose D, Backofen R. 2012. GraphClust: alignment-free structural clustering of local RNA secondary structures. Bioinforma Oxf Engl 28: i224–232.

Hiller M, Pudimat R, Busch A, Backofen R. 2006. Using RNA secondary structures to guide sequence motif finding towards single-stranded regions. Nucleic Acids Res 34: e117.

Hofacker IL. 2009. RNA secondary structure analysis using the Vienna RNA package. Curr Protoc Bioinforma Chapter 12: Unit12.2.

Hogan DJ, Riordan DP, Gerber AP, Herschlag D, Brown PO. 2008. Diverse RNA-binding proteins interact with functionally related sets of RNAs, suggesting an extensive regulatory system. PLoS Biol 6: e255.

Kazan H, Ray D, Chan ET, Hughes TR, Morris Q. 2010. RNAcontext: a new method for learning the sequence and structure binding preferences of RNA-binding proteins. PLoS Comput Biol 6: e1000832.

Kerpedjiev P, Honer Zu Siederdissen C, Hofacker IL. 2015. Predicting RNA 3D structure using a coarse-grain helix-centered model. RNA N Y N 21: 1110–1121.

Kim D, Langmead B, Salzberg SL. 2015. HISAT: a fast spliced aligner with low memory requirements. Nat Methods 12: 357–360.

Lambert N, Robertson A, Jangi M, McGeary S, Sharp PA, Burge CB. 2014. RNA Bind-n-Seq: quantitative assessment of the sequence and structural binding specificity of RNA binding proteins. Mol Cell 54: 887–900.

Livi CM, Blanzieri E. 2014. Protein-specific prediction of mRNA binding using RNA sequences, binding motifs and predicted secondary structures. BMC Bioinformatics 15: 123.

Lukong KE, Chang K, Khandjian EW, Richard S. 2008. RNA-binding proteins in human genetic disease. Trends Genet TIG 24: 416–425.

Magoc T, Salzberg SL. 2011. FLASH: fast length adjustment of short reads to improve genome assemblies. Bioinforma Oxf Engl 27: 2957–2963.

Maticzka D, Lange SJ, Costa F, Backofen R. 2014. GraphProt: modeling binding preferences of RNA-binding proteins. Genome Biol 15: R17.

Nussinov R, Jacobson AB. 1980. Fast algorithm for predicting the secondary structure of single-stranded RNA. Proc Natl Acad Sci U S A 77: 6309–6313.

Pan X, Shen H-B. 2017. RNA-protein binding motifs mining with a new hybrid deep learning based cross-domain knowledge integration approach. BMC Bioinformatics 18: 136.

Rodriguez A, Laio A. 2014. Machine learning. Clustering by fast search and find of density peaks. Science 344: 1492–1496.

Sanford JR, Wang X, Mort M, Vanduyn N, Cooper DN, Mooney SD, Edenberg HJ, Liu Y. 2009. Splicing factor SFRS1 recognizes a functionally diverse landscape of RNA transcripts. Genome Res 19: 381–394.

Siddharthan R, Siggia ED, van Nimwegen E. 2005. PhyloGibbs: a Gibbs sampling motif finder that incorporates phylogeny. PLoS Comput Biol 1: e67.

Song Z, Wu P, Ji P, Zhang J, Gong Q, Wu J, Shi Y. 2012. Solution structure of the second RRM domain of RBM5 and its unusual binding characters for different RNA targets. Biochemistry (Mosc) 51: 6667–6678.

Toyoda H, Franco D, Fujita K, Paul AV, Wimmer E. 2007. Replication of poliovirus requires binding of the poly(rc) binding protein to the cloverleaf as well as to the adjacent C-rich spacer sequence between the cloverleaf and the internal ribosomal entry site. J Virol 81: 10017–10028.

Van Nostrand EL, Pratt GA, Shishkin AA, Gelboin-Burkhart C, Fang MY, Sundararaman B, Blue SM, Nguyen TB, Surka C, Elkins K, et al. 2016. Robust transcriptome-wide discovery of RNA-binding protein binding sites with enhanced CLIP (eCLIP). Nat Methods 13: 508–514.

Wang X, Juan L, Lv J, Wang K, Sanford JR, Liu Y. 2011. Predicting sequence and structural specificities of RNA binding regions recognized by splicing factor SRSF1. BMC Genomics 12 Suppl 5: S8.

Weyn-Vanhentenryck SM, Zhang C. 2016. mCarts: Genome-Wide Prediction of Clustered Sequence Motifs as Binding Sites for. Methods Mol Biol Clifton NJ 1421: 215–226.

Wilbert ML, Huelga SC, Kapeli K, Stark TJ, Liang TY, Chen SX, Yan BY, Nathanson JL, Hutt KR, Lovci MT, et al. 2012. LIN28 binds messenger RNAs at GGAGA motifs and regulates splicing factor abundance. Mol Cell 48: 195–206.

Wurth L. 2012. Versatility of RNA-Binding Proteins in Cancer. Comp Funct Genomics 2012: 178525.

Yeo GW, Coufal NG, Liang TY, Peng GE, Fu X-D, Gage FH. 2009. An RNA code for the FOX2 splicing regulator revealed by mapping RNA-protein interactions in stem cells. Nat Struct Mol Biol 16: 130–137.

Yuan X, Davydova N, Conte MR, Curry S, Matthews S. 2002. Chemical shift mapping of RNA interactions with the polypyrimidine tract binding protein. Nucleic Acids Res 30: 456–462.

Zarnack K, Konig J, Tajnik M, Martincorena I, Eustermann S, Stevant I, Reyes A, Anders S, Luscombe NM, Ule J. 2013. Direct competition between hnRNP C and U2AF65 protects the transcriptome from the exonization of Alu elements. Cell 152: 453–466.

Zhang C, Darnell RB. 2011. Mapping in vivo protein-RNA interactions at single-nucleotide resolution from. Nat Biotechnol 29: 607–614.

Zhang C, Lee K-Y, Swanson MS, Darnell RB. 2013. Prediction of clustered RNA-binding protein motif sites in the mammalian genome. Nucleic Acids Res 41: 6793–6807.

Zhang S, Zhou J, Hu H, Gong H, Chen L, Cheng C, Zeng J. 2016. A deep learning framework for modeling structural features of RNA-binding protein targets. Nucleic Acids Res 44: e32.

